# Multiplexed single-cell lineage tracing of mitotic kinesin inhibitor resistance in glioblastoma

**DOI:** 10.1101/2023.09.09.557001

**Authors:** Yim Ling Cheng, Matei A. Banu, Wenting Zhao, Steven S. Rosenfeld, Peter Canoll, Peter A. Sims

## Abstract

Glioblastoma (GBM) is a deadly brain tumor, and the kinesin motor KIF11 is an attractive therapeutic target because of its dual roles in proliferation and invasion. The clinical utility of KIF11 inhibitors has been limited by drug resistance, which has mainly been studied in animal models. We used multiplexed lineage tracing barcodes and scRNA-seq to analyze drug resistance time courses for patient-derived GBM neurospheres treated with ispinesib, a potent KIF11 inhibitor. Similar to GBM progression in patients, untreated cells lost their neural lineage identity and transitioned to a mesenchymal phenotype, which is associated with poor prognosis. In contrast, cells subjected to long-term ispinesib treatment exhibited a proneural phenotype. We generated patient-derived xenografts to show that ispinesib-resistant cells form less aggressive tumors *in vivo*, even in the absence of drug. Finally, we used lineage barcodes to nominate drug combination targets by retrospective analysis of ispinesib-resistant clones in the drug-naïve setting and identified drugs that are synergistic with ispinesib.

## INTRODUCTION

Failure of effective cancer treatment is due, in part, to the dynamic and evolving nature of tumors. Glioblastoma (GBM) is an incurable malignancy with a 6.9% five-year survival rate^1^; it is a highly heterogeneous and plastic tumor that is able to traverse multiple cell states in response to extrinsic and intrinsic cues^2–4^. Molecular profiling of GBM with single-cell RNA-seq (scRNA-seq) and other methods have identified co-occurring cellular states that were observed across patients along with transitions between them^2,5,6^. These states are likely differentially sensitive to treatment and can result in therapy-resistant populations that give rise to aggressive recurrent tumors.

Drug resistance can arise from both genetic selection and both pre-existing or adaptive transcriptional states, which lead to an advantageous phenotype^7–12^. Only limited genetic alterations are associated with recurrence in GBM, potentially indicating a larger role of non-genetic mechanisms in GBM^13–15^. Devising effective therapies requires tools for characterizing and identifying vulnerabilities in drug-resistant states.

Inhibitors of the kinesin-5 (also known as Eg5 and KIF11), a mitotic kinesin, are promising candidates for GBM therapy, but can be rendered ineffective due to various mechanisms of resistance^16^. Kinesin-5 is a microtubule-associated motor protein that organizes the bipolar mitotic spindle and facilitates cell migration^17,18^. Its involvement in two hallmarks of cancer, proliferation and invasion, makes it a potential therapeutic target. GBM is deadly not only because GBM cells proliferate, but also because they invade the brain microenvironment. For inhibitors to be suitable for systemic GBM therapy, they also need to exhibit low neurotoxicity and high blood brain barrier penetration. Ispinesib is one such potent inhibitor and has been shown to prolong the survival of murine GBM models^19,20^. Resistance to this class of inhibitors can occur through point mutations that block drug binding^21^, upregulation of an alternative kinesin^22^, upregulation of drug efflux transporters^20^, upregulation of EGF to promote cell cycle progression^23^, and activation of STAT3 to inhibit apoptosis^24^.

To study the dynamics of GBM resistance and identify potential drug combination targets, we transfected PDGFR-amplified, patient-derived glioma neurospheres (TS543^25^) with a barcoded lineage tracing library (CellTag^26^), and treated the neurospheres with ispinesib. These genetically-modified, patient-derived neurospheres recapitulate key aspects of GBM heterogeneity and allow for surveillance of resistant phenotypes on multiple timescales. Lineage tracing barcodes allow us to selectively analyze clones that are destined for resistance in the drug-naïve setting. We analyzed the phenotypes of the glioma cells during the long-term ispinesib treatment with single-cell RNA-seq (scRNA-seq), assessed the stability and survival impact of drug-resistant phenotypes in the absence of drug and in orthotopic xenografts, and identified molecular markers of resistant clones in the drug-naïve setting to nominate effective drug combinations.

## RESULTS

### The kinesin-5 inhibitor, ispinesib, prevents mesenchymal transformation, resulting in a proneural resistant population

We first sought to compare the phenotypes of glioma cells after long-term treatment with ispinesib and DMSO (vehicle). We performed long-term treatment experiments (Figure 1A) as follows: 1) small replicate populations of TS543 were seeded and either transfected with CellTag^27^, a lentivirus barcode library, (celltag1 and celltag2) or not (control1 and control2); 2) the cells were expanded prior to treatment; 3) scRNA-seq was performed on the drug naïve cells; 4) after drug treatment started, scRNA-seq was initially performed at three-day intervals, and later at two- and four-week intervals; 5) viability of cells was monitored (Figure 1B) during this period to monitor drug resistance.

**Figure 1.**
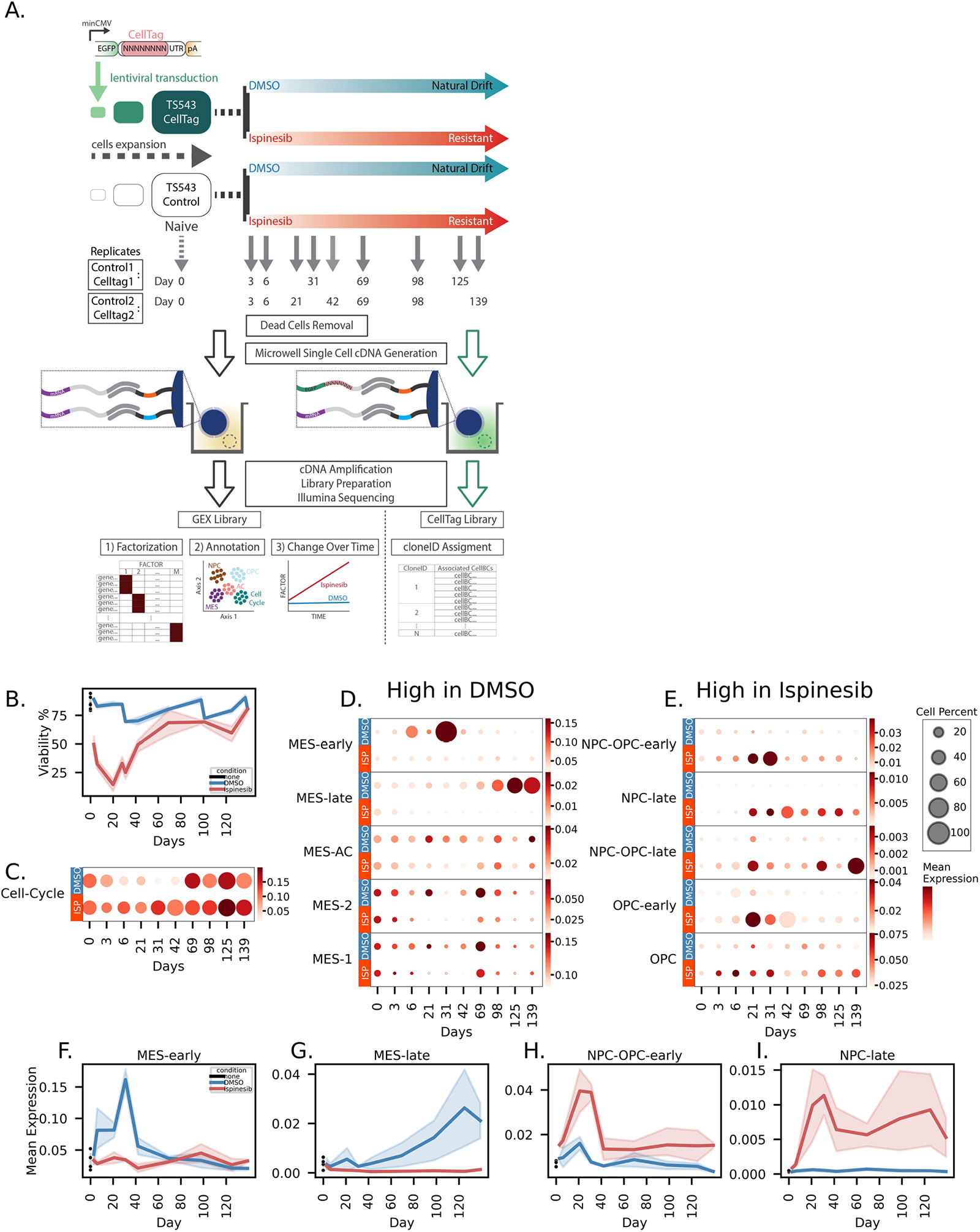
Ispinesib prevents mesenchymal transformation, and resistant population harbors proneural phenotype. A) Experimental schematic of single-cell lineage tracing and ispinesib-resistance time course in TS543 glioma neurospheres. B) Viability of TS543 during treatment. C-E) Dot plots of cell cycle factor (C), MES/AC associated factors (D), and NPC/OPC associated factors (E), with size indicating the percentage of cells with high cell-scores for the factor and color gradient indicating the mean log-normalized gene expression of top-ranked genes of the factor. F-I) Line plots of mean log-normalized gene expression of top-ranked genes in MES-early factor (F), MES-late factor (G), NPC-OPC-early factor (H), and NPC-late factor (I).

To identify cellular states and temporal patterns in the resulting dataset, we first pooled scRNA-seq data from all four replicates (control1, control2, celltag1, celltag2) and all time points to construct a factor model using single-cell hierarchical Poisson factorization (scHPF)^28^, a probabilistic algorithm for identifying co-expression signatures from scRNA-seq. scHPF computed scores that rank the association of each gene (Figure S1A, Supplementary Table 1) and cell with each identified co-expression signature or factor. We noticed that the highly ranked genes in many of the factors correspond to gene signatures reported in previous single-cell studies of GBM^2^. Thus, we systematically identified factors with high gene-scores for the cellular subtype gene sets from Neftel et al (Figure S1B). These gene sets were derived from the integration of scRNA-seq from 28 GBM and 401 bulk RNA-seq profiles from The Cancer Genome Atlas (TCGA)^2^ and correspond to cell cycle (G1/S, G2/M), hypoxic and non-hypoxic mesenchymal-like (MES1, MES2 respectively), astrocytic-like (AC), oligodendrocyte precursor-like geneset (OPC), and neural-progenitor-like (NPC1, NPC2)^2^. For factors with a clear temporal pattern, we appended the term “early” to the factor annotation, if the factor has high expression mainly before day 42, or “late”, if the factor has high expression mainly after day 42. In total, we annotated 11 factors: Cell-Cycle-1, Cell-Cycle-2, NPC-late, NPC-OPC-late, NPC-OPC-early, OPC-early, OPC, MES-early, MES-1, MES-2, MES-late, MES-AC (Figure S1B, Figure 1C-1E); top-ranked genes of Cell-Cycle-1 and Cell-Cycle-2 were combined into a single cell cycle factor.

Across our time course, we observed a clear enrichment of MES-associated factors in the DMSO samples (Figure 1D) and NPC-/OPC-associated (proneural) factors in the ispinesib-treated samples (Figure 1E). Specifically, we observed emergence of the MES-early cell state followed by a gradual increase in the MES-late cell state in the DMSO time course (Figure 1D, 1F, 1G). In contrast, we observed an early emergence in NPC-OPC-early and OPC-early cell states followed by an increase of NPC-late and NPC-OPC-late cell states for ispinesib-treated samples (Figure 1E, 1H, 1I). These results were reproducible among the four replicates (Figure S2A-S2B). To visualize the similarity of the four replicates, we also embedded the DMSO and ispinesib samples separately into a low-dimensional space with PHATE^29^. PHATE embeddings for the DMSO and ispinesib datasets were also reproducible across the four replicates (Figure S3A-S3D). The dominant temporal patterns are drift of the DMSO-and ispinesib-treated cells towards MES and NPC states, respectively.

The drift toward a mesenchymal phenotype is common phenomenon in solid tumors including GBM^30–34^ where it is associated with recurrence, drug resistance, and poor prognosis^35^. While it is unsurprising to observe mesenchymal drift in the DMSO time course as described above, this does not occur in the ispinesib-treated populations. Thus, although glioma cells develop resistance to ispinesib, the resistant cells resemble the proneural glioma phenotype, which is associated with improved survival^36^.

### The proneural phenotype of ispinesib-resistant clones is preserved in drug-free xenograft and associated with better survival

Our scRNA-seq time course revealed that ispinesib-resistant cells become increasingly proneural over time, whereas DMSO cells drift towards a mesenchymal phenotype associated with recurrence and poor survival^37,38^. Thus, we hypothesized that, unlike drug-resistant cells in many other settings, ispinesib-resistant cells might form less aggressive tumors than the untreated population. To test this hypothesis, we generated xenograft models by orthotopic transplantation of ispinesib-resistant and DMSO-natural-drift cells from last *in vitro* time point (day 125 control1 samples) (Figure 2A). Importantly, these xenografts were formed in the absence of drug. As expected, xenografted mice derived from ispinesib-resistant cells have a significant survival advantage (median survival 33 and 47 days for xenografts from DMSO-and ispinesib-treated cells, respectively; p = 8x10^-6^, log-rank test; Figure 2B), with no mice injected with ispinesib-resistant cells dying before any of the mice injected with DMSO-natural-drift cells.

**Figure 2.**
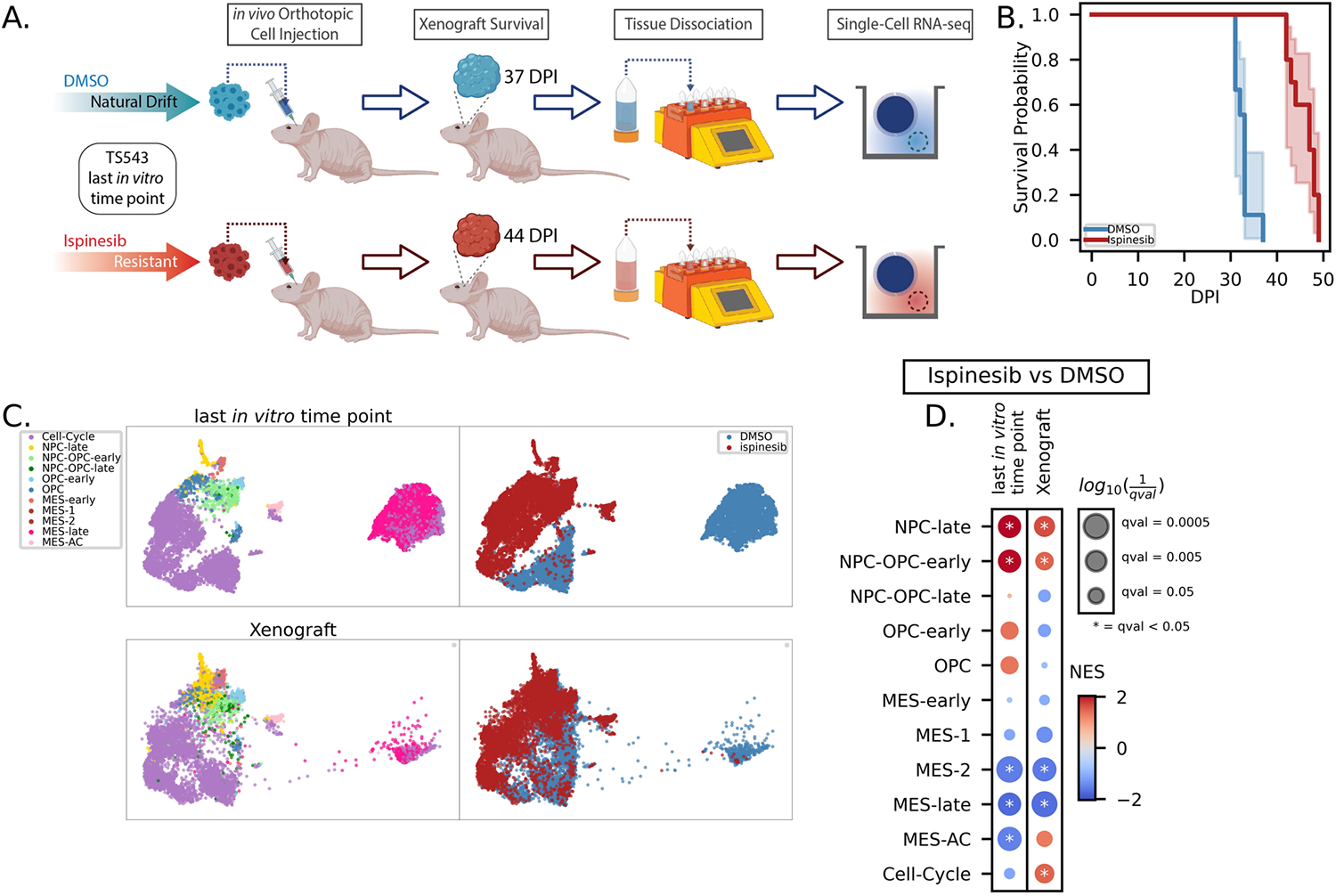
Proneural phenotype of ispinesib-resistant clones is preserved in drug-free xenograft and provides survival advantage. A) Experimental schematic of scRNA-seq of xenografts derived from DMSO-treated and ispinesib-resistant clones of last *in vitro* time point (day 125 control1 samples). B) Kaplan-Meier survival curves of ispinesib-resistant and DMSO-natural-drift clones derived xenograft models (p = 8x10^-6^, log-rank test). C) UMAP embeddings of last *in vitro* time point scHPF factors and xenograft dataset projection onto the scHPF model of the *in vitro* time course, color-coded by cell-states and treatments. D) Preranked GSEA normalized enrichment score (NES) of gene sets created from scHPF top-ranked genes and pre-ranked gene lists generated from differential expression analysis results between ispinesib and DMSO samples.

Given that these xenografts formed over the course of several weeks in the brain microenvironment and in the absence of drug, we next asked whether the phenotypic differences between the DMSO-natural-drift and ispinesib-resistant cells were preserved in end-stage tumors. We performed scRNA-seq on end-stage tumors from each population. To compare the phenotypes between the end-stage tumors of xenografts and the last *in vitro* time point which these xenografts were derived, we projected the scRNA-seq data from xenografts into the scHPF model of the original *in vitro* time course and visualized the xenograft projection along with the last *in vitro* time point with UMAP embedding (Figure 2C). The ispinesib-resistant and DMSO xenograft profiles project onto the *in vitro* ispinesib-resistant and DMSO-natural-drift populations, respectively, suggesting that key aspects of the ispinesib-resistant and DMSO-natural-drift phenotypes are preserved *in vivo* in the absence of treatment (Figure 2C).

To confirm that the mesenchymal phenotype of DMSO-natural-drift cells and proneural phenotype of ispinesib-resistant cells were maintained in the xenograft, we compared the two populations using differential expression analysis. We then used GSEA to identify scHPF signatures that are statistically enriched among the differentially expressed genes (Figure 2D). NPC-OPC-early and NPC-late signatures of ispinesib-resistant cells and MES-2 and MES-late signatures of DMSO-natural-drift cells are maintained in the xenograft (Figure 2D). Thus, despite growing in the absence of ispinesib for more than a month, ispinesib-resistant cells maintain their mesenchymal-depleted, proneural-enriched phenotype, consistent with the observed enhanced survival.

### Ispinesib-resistant clones are phenotypically diverse in the drug-naïve setting

The CellTag barcodes allow us to retrospectively analyze the phenotype of the resistant clones in the drug-naïve setting and determine whether they have unique properties relative to the remaining cells. To characterize the cell-states of clones that would become resistant, we partitioned the naïve samples into detected-future-resistant clones and remaining clones by matching the cloneIDs (assigned IDs of the multiplexed CellTag barcodes in clones) in the naïve samples with the cloneIDs in the ispinesib-exposed samples (Figure 3A). We used the term “remaining” instead of the term “sensitive” to name the naïve clones that did not have matched cloneIDs in ispinesib-exposed samples because of the possibility that not all ispinesib-exposed resistant clones were sequenced due to limited cell sampling. We embedded the scHPF factors of naïve samples in two-dimensions with UMAP to visualize any bias in the cell states occupied by the detected-future-resistant clones. In the UMAP embedding, we have the proliferating cells on the left side (Figure 3B). The quiescent population on the right side includes cells enriched in the OPC-and NPC-like signatures (Figure 3E-3G) with more mesenchymal (MES-early) cells on the far right (Figure 3C-3D). The detected-future-resistant clones are distributed throughout the UMAP with no visually obvious bias in the distribution of cell states for the detected-future-resistant clones (Figure 3H, 3I). To analyze this more quantitatively and determine if there are any significant cell-state differences between detected-future-resistant clones and remaining clones, we used GSEA to identify scHPF factors with enrichment among the genes that were differentially expressed between the detected-future-resistant clones and remaining clones (Figure 3J). Depletion of MES-early factor is significant for both replicates (celltag1, celltag2) (Figure 3J). While the other factors do not exhibit significant enrichment or depletion in the resistant clones, many of them have reproducible patterns such as enrichment of NPC-late, NPC-OPC-early and Cell-Cycle factors and depletion of MES-late and MES-1 factors (Figure 3J).

**Figure 3.**
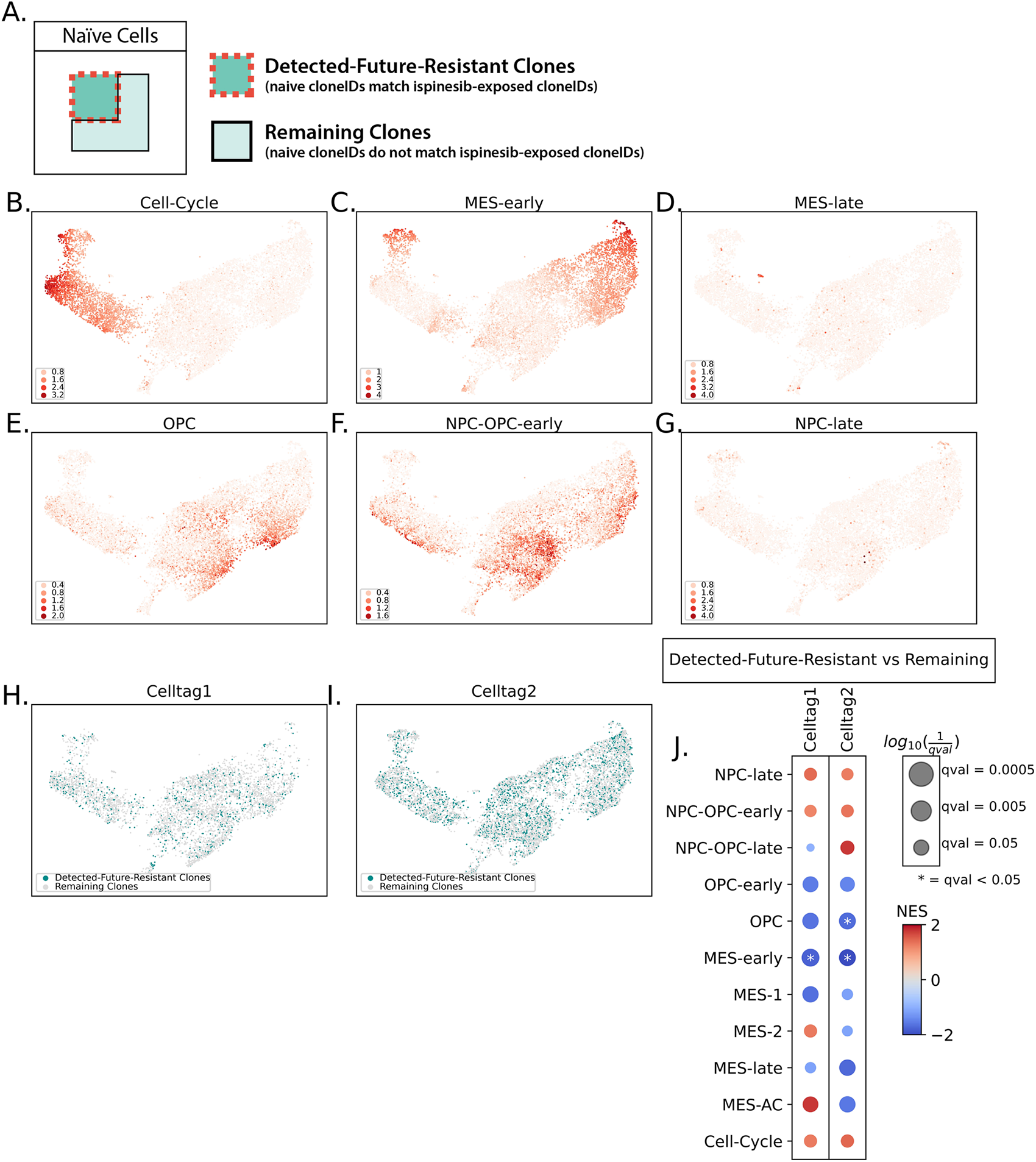
Ispinesib-resistant clones are phenotypically diverse in the naïve setting. A) Definitions of detected-future-resistant clones and remaining clones. CloneIDs are the assigned IDs of the multiplexed CellTag barcodes in clones. B-G) UMAP embedding of scHPF factors of naïve cells, colored by the cell-scores of cell-cycle factor (B), MES-early factor (C), MES-late factor (D), OPC factor (E), NPC-OPC-early factor (F), and NPC-late factor (G). H-I) Detected-future-resistant clones in celltag1 and celltag2 naïve cells on the UMAP embedding. J) Preranked GSEA normalized enrichment score (NES) of gene sets created from scHPF top-ranked genes and pre-ranked gene lists generated from differential expression analysis results between detected-future-resistant clones and remaining clones.

While the cells exhibit a strong phenotype after long-term treatment, there is no dominant phenotype for the detected-future-resistant clones in the naïve setting, and only subtle phenotypic differences between detected-future-resistant and remaining clones. This could be because the treatment-naïve cells are highly plastic with cells transitioning between these states on relatively short timescales or because acquisition of ispinesib-resistance states is stochastic. Nonetheless, the significant and reproducible depletion of the early mesenchymal (MES-early) state in the resistant clones is consistent with the depletion of mesenchymal expression signatures in the resistant cells observed at later time points.

### Identification of synergistic drug targets from gene expression analysis of ispinesib-resistant clones in the drug-naïve setting

We next sought to use our clonal lineage tracing data to identify druggable markers of resistant clones found in drug-naïve cells. By targeting druggable markers of clones that become resistant to ispinesib, we might identify effective therapies to be used in combination with ispinesib. Treatment time was divided into three periods: early (6 days and earlier), middle (21 to 42 days), and late (69 days or later) (Figure 4A). CloneIDs of each period were used to identify clones in naïve cells (Figure 4A). For each time period, we performed differential expression analysis between detected-future-resistant clones and remaining clones among drug-naïve cells (Figure 4B). Differentially expressed genes were ranked by fold-change and consistency between replicates (Figure 4C). Genes were removed if they were: 1) also differentially expressed in clones that were selected by DMSO or 2) encoding a protein for which a commercial inhibitor was unavailable (Figure 4C). We further identified genes with increasing expression during our time course (Figure 4D). Ultimately, we identified nine gene targets with available inhibitors that we could test in combination with ispinesib.

**Figure 4.**
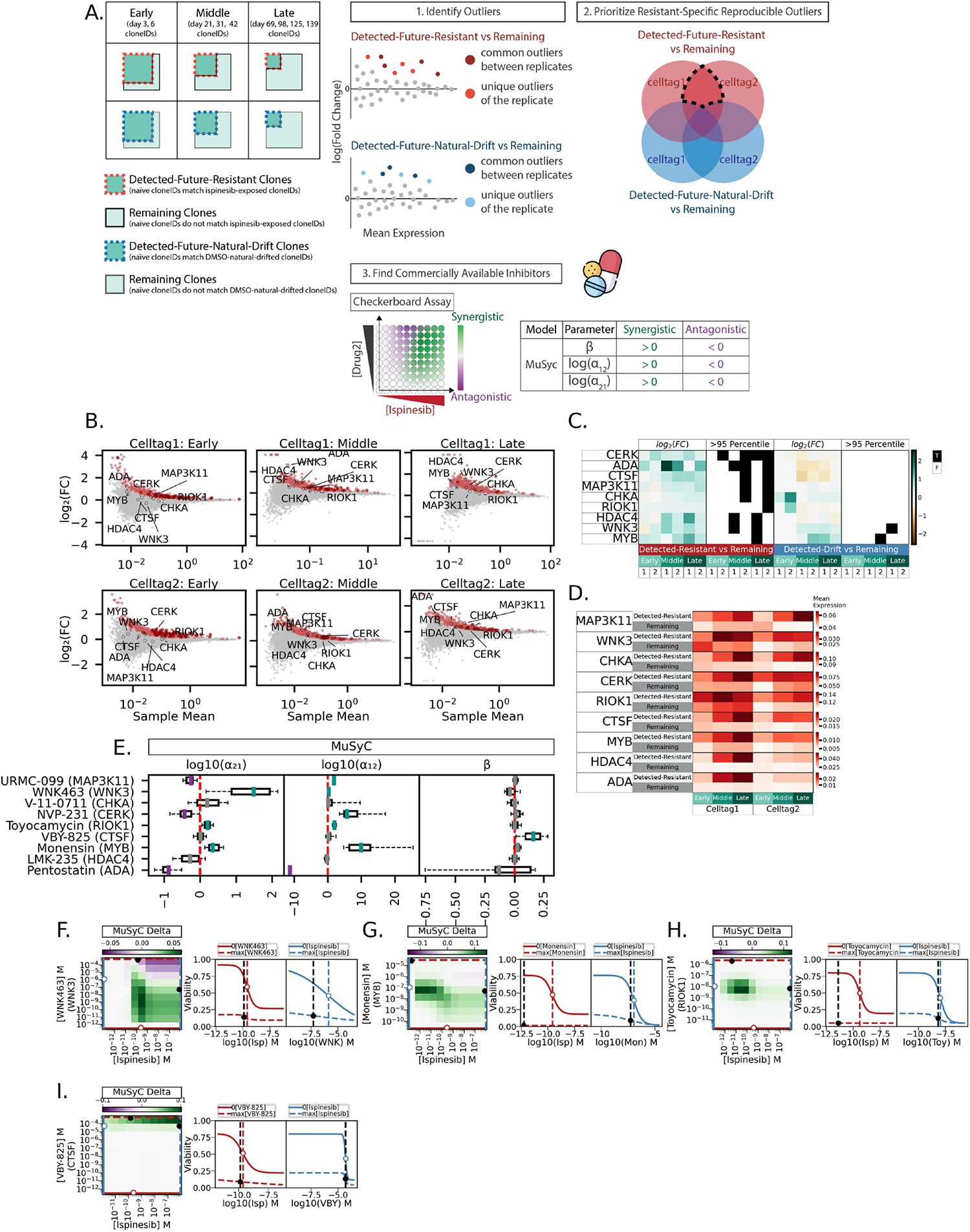
Identification of synergistic drug targets from gene expression analysis of ispinesib-resistant clones in the drug-naïve setting. A) Analysis scheme for identifying potential drug combinations with ispinesib. B) MA plots of log_2_(fold change) and sample mean from the differential expression analysis between detected-future-resistant clones and remaining clones of naïve cells. Genes with fold changes that were significantly high (>95 percentile) given their sample means were colored as red. C) Heatmaps of log_2_(fold change) from the differential expression analysis between detected clones and remaining clones of naïve cells. Boolean heatmaps of whether the log_2_(fold change) was significantly high (>95 percentile) given the genes’ sample means. In the x-axis labels of heatmaps, 1 is celltag1 sample, 2 is celltag2 sample, detected-resistant is the alias for detected-future-resistant clones, and detected-drift is the alias for detected-future-natural-drift clones. D) Mean log-normalized gene expression of the druggable target genes of detected-future-resistant and remaining clones of naïve cells. E) Boxplots of synergistic parameters of MuSyC models. F-I) Heatmaps of MuSyc Delta, which is difference between the 2D Hill-fitted model of the observed efficacy and the null hypothesis. Side-by-side single drug response curve (solid curve line, with the other drug concentration at zero) and combined drugs response curve (dash curve line, with the other drug concentration at the maximum tested).

We performed a drug interaction experiment with the “checkerboard” assay and analyzed the results with MuSyc (multi-dimensional synergy of combinations)^39^, a recently reported multi-drug synergy framework. MuSyc is a two-dimensional extension of the traditional single drug Hill equation and has five additional parameters that quantify drug interactions: efficacy (β), potency (⍺_12_, ⍺_21_), and cooperativity (ɣ_12_, ɣ_21_). The intention of drug combination treatment is to reduce toxicity by minimizing dose, to improve survival by increasing efficacy, or both. The parameter ⍺ quantifies the change in effective dose (EC_50_) of the first drug in the presence of the second drug. For synergistic potency (⍺ > 1 or log(⍺) > 0), EC_50_ of the first drug decreases in the presence of the second drug, which corresponds to increase of potency. ⍺_21_ is the fold change in potency of drug1, which was ispinesib in our case, induced by the presence of drug2, and vice versa. The parameter β quantifies the change in the maximal effect of the two drugs in combination compared to the maximal effect of the most efficacious single drug. For synergistic efficacy (β > 0), the combined effect at the maximum concentration tested for both drugs is greater than the maximum effect of either drug alone.

WNK463, toyocamycin, and monensin, which are inhibitors of WNK3 (Lysine-deficient protein kinase 3), RIOK1 (RIO kinase 1), and MYB (MYB proto-oncogene transcription factor) respectively, have synergistic potencies when combined with ispinesib; log(⍺_12_) and log(⍺_21_) are above zero (Figure 4E). VBY-825, an inhibitor of CTSF (Cathepsin F), when combined with ispinesib has synergistic efficacy; β is greater than zero (Figure 4E). The heatmap of MuSyc Delta, which is the difference between the 2D Hill-fitted model of the observed efficacy and the null hypothesis, shows regions where combined dosage of two drugs have efficacy greater than the null hypothesis (Figure 4F-4I). The side-by-side dose-response curves show the single drug response curve (solid curve line, with the other drug concentration at zero) and combined drugs response curve (dash curve line, with the other drug concentration at the maximum tested). Ispinesib alone reaches maximum efficacy of 25% to 20% viability (red solid curve lines), and the viability is further decreased with addition of the second drug (blue dash curve lines) (Figure 4F-4I). WNK3^40–42^, RIOK1^43–45^, MYB^46,47^ and CTSF^48^ may protect ispinesib-resistant cells from apoptosis or help ispinesib resistant cells progress through cell cycle. Overall, these results demonstrate how lineage tracing analysis by scRNA-seq can be used to identify novel drug combinations by identifying markers of resistant clones in the drug-naïve setting.

## DISCUSSION

GBM often contains cells with a mesenchymal phenotype that is associated with poor survival and drug resistance and becomes more pronounced upon recurrence. Similarly, the glioma neurosphere model used here drifts towards a mesenchymal phenotype in the absence of treatment. We found that the kinesin-5 inhibitor, ispinesib, effectively prevents this mesenchymal transition, and the resistant population that emerges instead harbors a proneural phenotype. Thus, we reasoned that the ispinesib-resistant cells would form less aggressive tumors than the more mesenchymal cells observed in the absence of drug. While targeted therapies often select for more aggressive phenotypes, we found that the ispinesib-resistant clones formed significantly less aggressive orthotopic xenografts, even in the absence of drug. Subsequent scRNA-seq analysis confirmed that the phenotypic differences between the ispinesib-resistant and DMSO-treated cells were largely preserved in the animal model. These findings raise the exciting possibility that ispinesib could not only serve as an effective targeted therapy in GBM, but that the ispinesib-resistant clones that arise may be less aggressive than the mesenchymal cells typically found after standard treatment.

Despite the less aggressive phenotype of ispinesib-resistant cells, it remains desirable to identify drug combinations with the potential to minimize resistance. The single-cell lineage tracing approach used here provides the unique ability to retrospectively analyze the phenotype of ispinesib-resistant clones in a treatment-naïve population. This analysis identified genes that were enriched in drug-naïve clones that would become resistant to ispinesib, some of which encoded druggable protein targets. Perhaps not surprisingly, subsequent validation experiments showed that targets associated with cell survival and apoptosis such as WNK3, RIOK1, MYB, and CTSF were synergistic with ispinesib. Previous studies of ispinesib resistance by Kenchappa et al^24^ also concluded that glioma cells activate anti-apoptotic mechanisms to survive the prolonged G2M block produced by ispinesib, whereas normal cells apoptose under these conditions due to “mitotic catastrophe”. This phenomenon was shown to be mediated by STAT3 through its transcriptional activity and effects on mitochondrial membrane permeability and oxidative metabolism. Taken together, these studies show that multiple mediators of apoptosis could potentially be exploited by glioma cells to resist anti-mitotic drugs. Further efforts with long-term treatment and survival studies in animal models will be required to establish the pre-clinical efficacy of these interesting new drug combinations. Nonetheless, the strategy employed here for discovering these drug combinations has significant advantages over conventional combinatorial screening in rapidly narrowing the scope of potential candidates targeted to drug-resistant clones.

## METHODS

### Cell lines and cultures

PDGFRA-amplified, patient-derived glioblastoma neurospheres, TS543^25^, were cultured with NeuroCult^TM^ NS-A Proliferation Kit Human from STEMCELL Technologies. HEK293T were cultured with DMEM containing 10% FBS and 2 mM L-glutamine.

### CellTag barcode lentivirus transduction and long-term ispinesib treatment

TS543 cells were seeded at ∼1000 cells and were transduced with Celltag^27^ virus-laden media at MOI of around 8-10 and 5 ug/ml protamine sulfate or normal media and 5 ug/ml protamine sulfate. TS543 were propagated for three weeks before start of treatment. scRNA-seq was performed on drug-naïve TS543. TS543 were treated with 75 nM ispinesib or with vehicle DMSO; media with ispinesib or DMSO were replenished every two or three days. Viability was monitored as guidepost for resistance. Dead cells were removed with Dead Cell Removal Kit from Miltenyi Biotec; scRNA-seq was initially performed at three-day interval, and later at two-and four-week intervals during treatment.

### CellTag barcode lentivirus packaging

CellTag barcode lentivirus was packaged according to the online protocol on protocols.io^27^. Briefly, lentiviral pSMAL-CellTag-V1 pooled library and its associated packing plasmids pCMV-dR8.2 dvpr and pCMV-VSV-G were obtained from Addgene, lentiviruses were produced by transfecting with HEK293T cells using X-tremeGENE^TM^ 9 DNA Transfection Reagent from Sigma-Aldrich, and virus was collected 48 hours after transfection. Virus was concentrated with Lenti-X^TM^ Concentrator from Takara and re-suspended in NeuroCult^TM^ NS-A Complete Media.

### Animals

All procedures were reviewed and approved by the Columbia University Institutional Animal Care and Use committee (IACUC). Nude CrTac:NCr-Foxn1^nu^ female mice (Taconic Biosciences) were used as background for *in vivo* orthotopic cell injection experiments. Mice were housed in pathogen-free facilities at Columbia University Irving Medical Center. Mice were ordered and housed under standard conditions after arrival.

### Murine glioma models and survival analysis

For orthotopic cell transplantation experiments, six-week-old CrTac:NCr-Foxn1^nu^ female mice were injected with 5 x 10^4^ cells of last *in vitro* time point of TS543 treated with ispinesib or vehicle DMSO (day 125 control1 samples); ten mice were used per cohort. Mice were clinically monitored daily and sacrificed once end-stage criteria were met, including severe weight loss, seizures, and evidence of motor deficit. Tissues of one mouse from each cohort were harvested and processed for scRNA-seq. Survival curves were modeled by Kaplan-Meier method.

### Tissue dissociation

Mouse brain tumor resections were dissociated using Adult Brain Dissociation kit on gentleMACS^TM^ Octo Dissociator with Heaters (Miltenyi Biotec) according to manufacturer’s instructions.

### Microwell scRNA-seq

Microwell 3’ scRNA-seq was performed as described^49^. Briefly, individual cells were co-encapsulated with a barcoded mRNA capture bead^50^ (MACOSKO-2011-10, ChemGenes) and lysed in microwell-based platform, mRNA transcripts were captured and reverse transcribed on the bead, cDNA-coated beads were pooled for PCR amplification, and Illumina Nextera libraries were constructed for each sample. Gene expression libraries were sequenced on an Illumina NovaSeq 6000 with 51 cycles or 26 cycles for read 1 and 151 cycles for read 2.

To sequence the CellTag library separately from the gene expression library at greater sequencing depth, CellTag libraries were constructed with custom P5_TSO_hybrid primer^50^ and custom P7_TruSeq-6bp-Unique-Index_EGFP primer (CAAGCAGAAGACGGCATACGAGAT[6bp-RPI]GTGACTGGAGTTCCTTGGCACCCGAGAATTCCAGGCATGGACGAGCTGTACAAGT*A*A) from the barcoded cDNA libraries. CellTag libraries were sequenced on NextSeq 550 (Illumina) with 26 cycles for read 1 and 58 cycles for read 2.

### scRNA-seq preprocessing and quality control

Raw data obtained from NovaSeq was corrected for index swapping according to the BarcodeSwapping method^51^. Raw reads were preprocessed as described^5^, with a brief description as follows. Raw reads were subjected to polyA trimming and aligned with STAR. An address comprised of cell-barcode, UMI, and gene identifier was constructed for each read with a unique, strand-specific alignment to exonic sequence. Reads with same address were collapsed and sequencing errors in cell-barcodes and UMI were corrected. Cell-barcodes of empty microwell or low quality cells were removed. Empty cell-barcodes were identified with EmptyDrops algorithm^52^. Cells were filtered as low-quality if they meet any of the following criteria: 1) fractional alignment to the mitochondrial genome per cell-barcode is greater than 10%, 2) the ratio of molecules aligning to whole gene bodies (including introns) to molecules aligning exclusively to exons is greater than 1.96 standard deviations above the mean, 3) average number of reads per molecule or average number of molecules per gene is greater than 2.5 standard deviations above the mean, or 4) more than 40% of UMI bases are T or where the average number of T-bases per UMI is at least 4.

### CellTag processing and clone calling

CellTag binary count matrices were generated and clone callings were performed with the CellTagWorkflow algorithm (https://github.com/morris-lab/CellTagWorkflow), with an additional preprocessing step of removing UMI with less than three raw reads.

### Single-cell hierarchical Poisson factorization (scHPF)

Single-cell RNA-seq data from all four replicates (control1, control2, celltag1, celltag2) and all time points were pooled to construct a factor model using single-cell hierarchical Poisson factorization (scHPF)^28^. scHPF outputted a list of factors and factor-associated gene-scores for each gene (Figure S1A, Supplementary Table 1) and cell-scores for each cell of that factor, with higher score being more associated with the factor. Based on the top ranked genes in each factor, we removed factors with high gene-scores for ribosomal genes, which represented sequencing coverage, and high gene-scores for interferon genes, which was associated with lentivirus transfection, from downstream analysis. To annotate the factors, we looked for factors with high gene-scores for Neftel-glioblastoma gene sets^2^ (Figure S1B).

### Differential expression and GSEA

Count matrices for the two conditions were subsampled to give the same cell numbers and same average number of unique transcripts per cell. The resulting count matrix was normalized by the scran deconvolution approach^53^. Differential expression analysis between two conditions was performed using the Mann-Whitney U-test (*scipy.mannwhitneyu*). The p-values were adjusted for false discovery with Benjamini-Hochberg procedure (*statsmodels.multipletests*). Genes were ranked by log_2_(fold change) × -log_10_(FDR adjusted p-value). Preranked GSEA was performed on the ranked genes from differential expression analysis with gene sets created from the top scoring genes of scHPF factors.

### Dose response assay and drug synergy analysis

Single drug dose response assays were performed on each inhibitor to determine their IC50. TS543 cells were seeded in 96 wells plate at concentration of 1 x 10^4^ cells/cm^2^ and grown for four days and treated with inhibitor for three days. Cell viability was assessed with PrestoBlue^TM^ Cell Viability Reagent from ThermoFisher. Single drug dose response curves were fitted with the Hill model (*synergy.single.Hill*), and IC50 was determined for each inhibitor. With IC50 as the mid-range concentration, two-drug eight-by-eight dose response checkerboard assays were performed with ispinesib and a candidate inhibitor. TS543 cells were seeded in 384 wells plate at concentration of 1 x 10^4^ cells/cm^2^, grown for four days and treated with ispinesib, and candidate inhibitor for three days. Cell viability was assessed with PrestoBlue^TM^ Cell Viability Reagent from Thermo Fisher. Two-drug dose response surfaces were fit with the MuSyc model^39^ (*synergy.combination.MySyc*, https://github.com/djwooten/synergy), and synergistic parameters were determined for each drug combination. Ispinesib, URMC-099, WNK463, tomocamycin, NVP231, LMK-235, pentostatin, and monensin sodium salt were purchased from MedChemExpress. VBY-825 and V-11-0711 were purchased from AdooQ Bioscience and MedKoo Bioscience respectively.

## DATA AVAILABILITY

Raw and preprocessed data in this study are available in the NCBI Gene Expression Omnibus (GEO) with accession code GSE239651.

## AUTHOR CONTRIBUTIONS

Y.L.C. designed and executed the lineage-tracing, drug treatment, scRNA-seq, and drug response experiments and performed associated analyses with inputs from P.A.S., P.C., and S.S.R. P.A.S preprocessed the raw data from NovaSeq. M.A.B generated and monitored the xenograft models and performed the survival analysis. W.Z. dissociated the mouse brain tumor resections. Y.L.C. and P.A.S. wrote the manuscript. All authors edited, read, and approved the final manuscript.

## COMPETING INTERESTS

P.A.S. is listed as an inventor on patent applications and issued patents filed by Columbia University related to the microwell technology described here. P.A.S. receives patent royalties from Guardant Health. The remaining authors declare that they have no competing interests.

## FUNDING

P.A.S., P.C., and S.S.R. were funded by R01NS118513 from NIH/NINDS. P.A.S. and P.C. were funded by R01NS103473 from NIH/NINDS. P.A.S. was funded by U01CA168426 from NIH/NCI.

## Notes

https://www.ncbi.nlm.nih.gov/geo/query/acc.cgi?acc=GSE239651

## REFERENCES

1. 1. About Glioblastoma. National Brain Tumor Society https://braintumor.org/events/glioblastoma-awareness-day/about-glioblastoma/.

2. Neftel, C. et al. An Integrative Model of Cellular States, Plasticity, and Genetics for Glioblastoma. Cell 178, 835 (2019).

3. Al-Mayhani, T. M. F. et al. A non-hierarchical organization of tumorigenic ng2 cells in glioblastoma promoted by egfr. Neuro-oncology 21, 719–729 (2019).

4. Dirkse, A. et al. Stem cell-associated heterogeneity in Glioblastoma results from intrinsic tumor plasticity shaped by the microenvironment. Nature Communications 10, 1787–1787 (2019).

5. Yuan, J. et al. Single-cell transcriptome analysis of lineage diversity in high-grade glioma. Genome Medicine 10, 57 (2018).

6. Joshua S. Schiffman et al. Defining ancestry, heritability and plasticity of cellular phenotypes in somatic evolution. bioRxiv (2023) doi:10.1101/2022.12.28.522128.

7. Goyal, Y. et al. Diverse clonal fates emerge upon drug treatment of homogeneous cancer cells. Nature 1–9 (2023) doi:10.1038/s41586-023-06342-8.

8. Spencer, S. L., Gaudet, S., Albeck, J. G., Burke, J. M. & Sorger, P. K. Non-genetic origins of cell-to-cell variability in TRAIL-induced apoptosis. Nature 459, 428–432 (2009).

9. Wang, X. et al. Drug resistance and combating drug resistance in cancer. Cancer Drug Resistance 2, 141–160 (2019).

10. Bell, C. C. & Gilan, O. Principles and mechanisms of non-genetic resistance in cancer. British Journal of Cancer 122, 465–472 (2020).

11. Shen, S., Vagner, S. & Robert, C. Persistent Cancer Cells: The Deadly Survivors. Cell 183, 860–874 (2020).

12. Eyler, C. E. et al. Single-cell lineage analysis reveals genetic and epigenetic interplay in glioblastoma drug resistance. Genome Biology 21, 1–21 (2020).

13. Barthel, F. P. et al. Longitudinal molecular trajectories of diffuse glioma in adults. Nature 576, 112–120 (2019).

14. Körber, V. et al. Evolutionary Trajectories of IDHWT Glioblastomas Reveal a Common Path of Early Tumorigenesis Instigated Years ahead of Initial Diagnosis. Cancer Cell 35, 692 (2019).

15. Wang, J. et al. Clonal evolution of glioblastoma under therapy. Nat Genet 48, 768–776 (2016).

16. Garcia-Saez, I., Garcia-Saez, I. & Skoufias, D. A. Eg5 targeting agents: From new anti-mitotic based inhibitor discovery to cancer therapy and resistance. Biochemical Pharmacology 184, 114364 (2020).

17. Waitzman, J. S. & Rice, S. E. Mechanism and regulation of kinesin-5, an essential motor for the mitotic spindle. Biology of the Cell 106, 1–12 (2014).

18. Falnikar, A., Tole, S. & Baas, P. W. Kinesin-5, a mitotic microtubule-associated motor protein, modulates neuronal migration. Molecular Biology of the Cell 22, 1561–1574 (2011).

19. Venere, M. et al. The mitotic kinesin KIF11 is a driver of invasion, proliferation, and self-renewal in glioblastoma. Science Translational Medicine 7, (2015).

20. Gampa, G. et al. Enhancing Brain Retention of a KIF11 Inhibitor Significantly Improves its Efficacy in a Mouse Model of Glioblastoma. Scientific Reports 10, 6524–6524 (2020).

21. Talapatra, S. K., Anthony, N. G., Mackay, S. P. & Kozielski, F. Mitotic kinesin Eg5 overcomes inhibition to the phase I/II clinical candidate SB743921 by an allosteric resistance mechanism. Journal of Medicinal Chemistry 56, 6317–6329 (2013).

22. Sturgill, E. G., Norris, S. R., Guo, Y., Ohi, M. D. & Ohi, R. Kinesin-5 inhibitor resistance is driven by kinesin-12. Journal of Cell Biology 213, 213–227 (2016).

23. Mardin, B. R. et al. EGF-Induced Centrosome Separation Promotes Mitotic Progression and Cell Survival. Developmental Cell 25, 229–240 (2013).

24. Rajappa S Kenchappa et al. Activation of STAT3 through combined SRC and EGFR signaling drives resistance to a mitotic kinesin inhibitor in glioblastoma. Cell Reports 39, 110991–110991 (2022).

25. Silber, J. et al. miR-34a Repression in Proneural Malignant Gliomas Upregulates Expression of Its Target PDGFRA and Promotes Tumorigenesis. PLOS ONE 7, e33844 (2012).

26. Kong, W. et al. CellTagging: combinatorial indexing to simultaneously map lineage and identity at single-cell resolution. Nature Protocols 15, 750–772 (2020).

27. Biddy, B. A. Single-cell mapping of lineage and identity via CellTagging. (2018) doi:10.17504/protocols.io.vawe2fe.

28. Levitin, H. M. et al. De novo gene signature identification from single-cell RNA-seq with hierarchical Poisson factorization. Molecular Systems Biology 15, (2019).

29. Moon, K. R. et al. PHATE: A Dimensionality Reduction Method for Visualizing Trajectory Structures in High-Dimensional Biological Data. bioRxiv 120378 (2017) doi:10.1101/120378.

30. Lin Wang et al. A single-cell atlas of glioblastoma evolution under therapy reveals cell-intrinsic and cell-extrinsic therapeutic targets. Nature Cancer (2022) doi:10.1038/s43018-022-00475-x.

31. Krishna P.L. Bhat et al. Mesenchymal Differentiation Mediated by NF-κB Promotes Radiation Resistance in Glioblastoma. Cancer Cell 24, 331–346 (2013).

32. Piao, Y. et al. Acquired Resistance to Anti-VEGF Therapy in Glioblastoma Is Associated with a Mesenchymal Transition. Clinical Cancer Research 19, 4392–4403 (2013).

33. Halliday, J. et al. In vivo radiation response of proneural glioma characterized by protective p53 transcriptional program and proneural-mesenchymal shift. Proceedings of the National Academy of Sciences of the United States of America 111, 5248–5253 (2014).

34. Wang, Q. et al. Tumor Evolution of Glioma-Intrinsic Gene Expression Subtypes Associates with Immunological Changes in the Microenvironment. Cancer Cell 32, 152–152 (2017).

35. Kim, Y. et al. Perspective of mesenchymal transformation in glioblastoma. Acta neuropathologica communications 9, 50 (2021).

36. Steponaitis, G. & Tamasauskas, A. Mesenchymal and Proneural Subtypes of Glioblastoma Disclose Branching Based on GSC Associated Signature. International Journal of Molecular Sciences 22, 4964 (2021).

37. Gill, B. J. et al. MRI-localized biopsies reveal subtype-specific differences in molecular and cellular composition at the margins of glioblastoma. Proceedings of the National Academy of Sciences 111, 12550–12555 (2014).

38. Olar, A. & Aldape, K. D. Using the molecular classification of glioblastoma to inform personalized treatment. The Journal of Pathology 232, 165–177 (2014).

39. Wooten, D. J., Meyer, C. T., Lubbock, A. L. R., Quaranta, V. & Lopez, C. F. MuSyC is a consensus framework that unifies multi-drug synergy metrics for combinatorial drug discovery. Nature Communications 12, 4607–4607 (2021).

40. Zhang, Z. et al. LINGO-1 Receptor Promotes Neuronal Apoptosis by Inhibiting WNK3 Kinase Activity. Journal of Biological Chemistry 288, 12152–12160 (2013).

41. Wu, D. et al. Activated WNK3 induced by intracerebral hemorrhage deteriorates brain injury maybe via WNK3/SPAK/NKCC1 pathway. Experimental Neurology 332, 113386 (2020).

42. Zhu, J. et al. WNK3 promotes neuronal survival after traumatic brain injury in rats. Neuroscience (2021) doi:10.1016/j.neuroscience.2021.09.021.

43. Weinberg, F. et al. The Atypical Kinase RIOK1 Promotes Tumor Growth and Invasive Behavior. EBioMedicine 20, 79–97 (2017).

44. Huang, Z. et al. Elevated Expression of RIOK1 Is Correlated with Breast Cancer Hormone Receptor Status and Promotes Cancer Progression. Cancer Research and Treatment 52, 1067–1083 (2020).

45. Rong Wang et al. RIOK1 is associated with non-small cell lung cancer clinical characters and contributes to cancer progression. Journal of Cancer 13, 1289–1298 (2022).

46. Ramsay, R. G. & Gonda, T. J. MYB function in normal and cancer cells. Nature Reviews Cancer 8, 523–534 (2008).

47. Cicirò, Y. & Sala, A. MYB oncoproteins: emerging players and potential therapeutic targets in human cancer. Oncogenesis 10, 19–19 (2021).

48. Yadati, T., Houben, T., Bitorina, A. V. & Shiri-Sverdlov, R. The Ins and Outs of Cathepsins: Physiological Function and Role in Disease Management. Cells 9, 1679 (2020).

49. Yuan, J. & Sims, P. A. An Automated Microwell Platform for Large-Scale Single Cell RNA-Seq. Sci Rep 6, 33883 (2016).

50. Macosko, E. Z. et al. Highly Parallel Genome-wide Expression Profiling of Individual Cells Using Nanoliter Droplets. Cell 161, 1202–1214 (2015).

51. Griffiths, J. A., Richard, A. C., Bach, K., Lun, A. T. L. & Marioni, J. C. Detection and removal of barcode swapping in single-cell RNA-seq data. Nat Commun 9, 2667 (2018).

52. Lun, A. T. L. et al. EmptyDrops: distinguishing cells from empty droplets in droplet-based single-cell RNA sequencing data. Genome Biology 20, 63 (2019).

53. L. Lun, A. T., Bach, K. & Marioni, J. C. Pooling across cells to normalize single-cell RNA sequencing data with many zero counts. Genome Biology 17, 75 (2016).

