## Supplementary Figures 1-3 for "Multiplexed single-cell lineage tracing of mitotic kinesin inhibitor resistance in glioblastoma"

**Supplementary Materials for: Multiplexed single-cell lineage tracing of mitotic kinesin inhibitor resistance in glioblastoma**

### SUPPLEMENTARY FIGURES

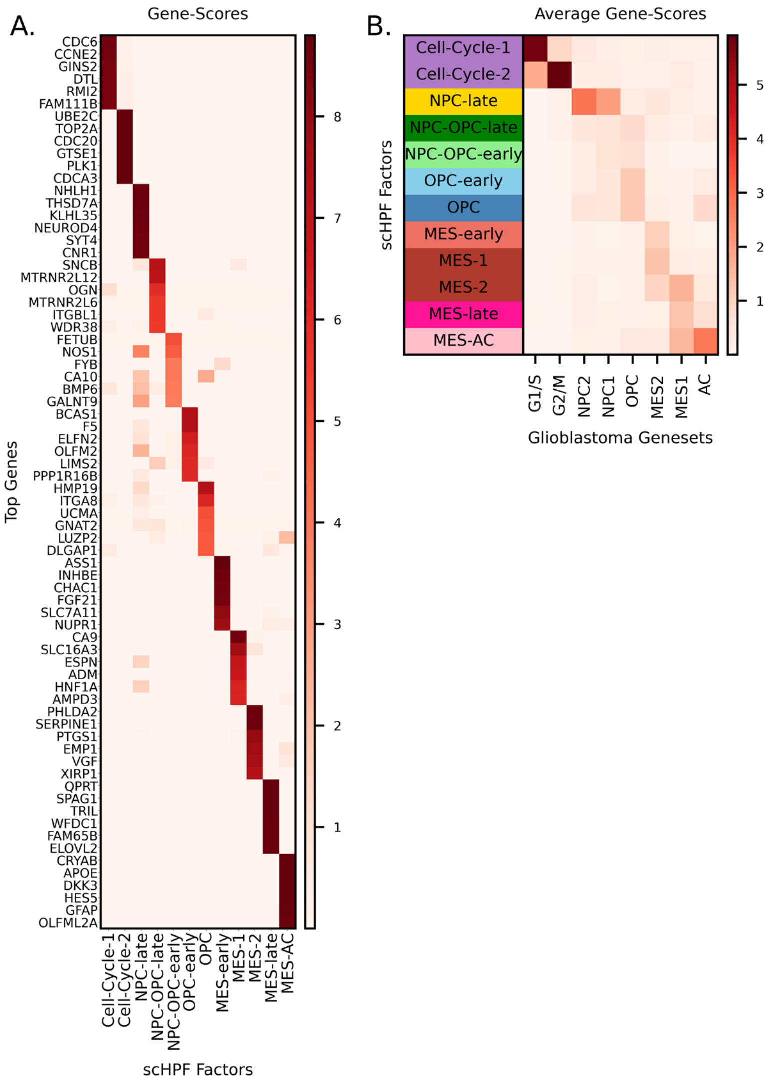


**Supplementary Figure 1. scHPF model of the pooled scRNA-seq data for the *in vitro* time course.** A) Heatmap of gene-scores of top-ranked genes of scHPF factors. B) Mean scHPF factor gene-scores for Neftel-glioblastoma gene sets.

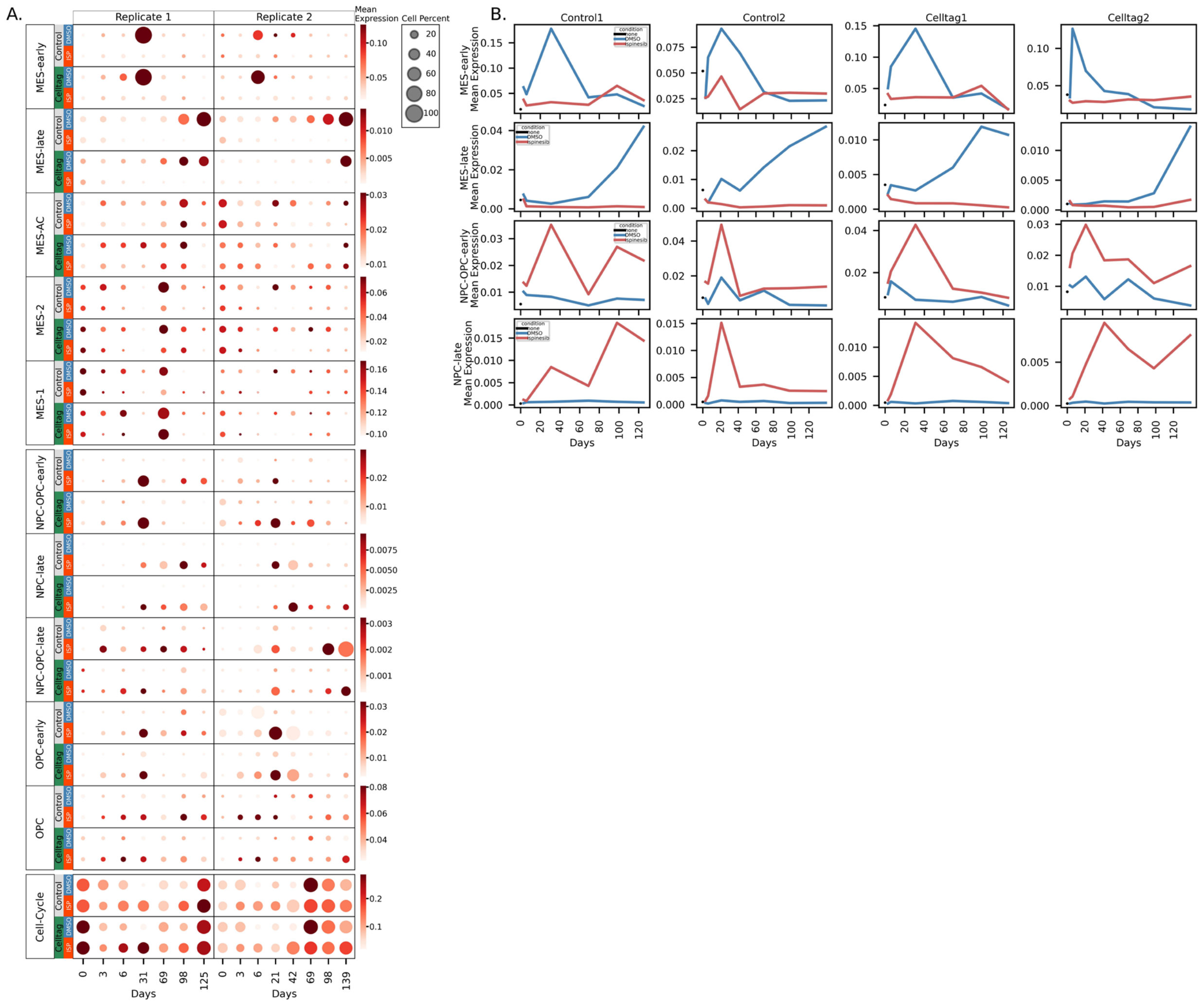


**Supplementary Figure 2. Phenotypic changes for DMSO and ispinesib-treated time course datasets were reproducible across four replicates.** A) Dot plots of scHPF factors with size indicating the percentage of cells with high cell-scores for the factor and color gradient indicating the mean log-normalized gene expression of top-ranked genes of the factor. B) Line plots of mean log normalized gene expression of top-ranked genes in MES-early factor, MES-late factor, NPC-OPC-early factor, and NPC-late factor.

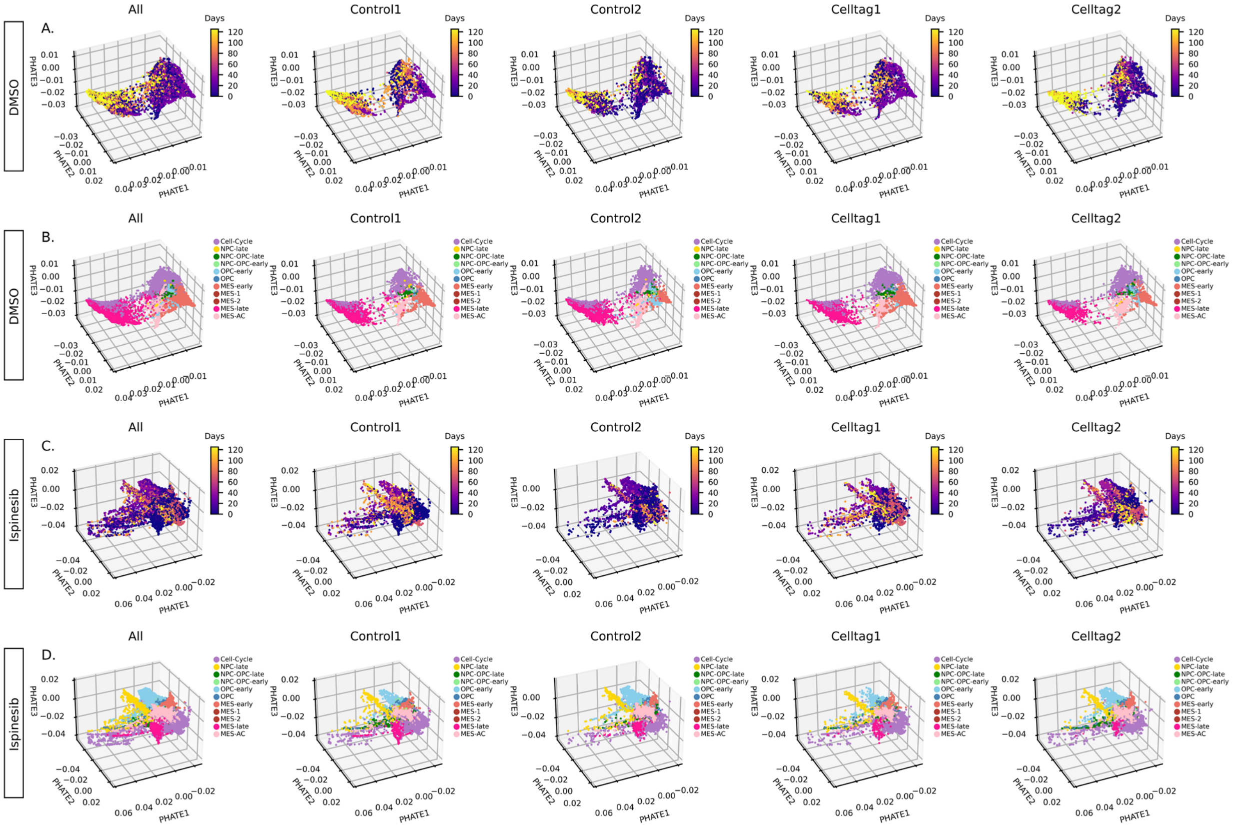


**Supplementary Figure 3. Trajectories of DMSO and ispinesib-treated time course datasets were reproducible across four replicates.** PHATE embeddings of DMSO samples' scHPF factors (A-B) and ispinesib samples' scHPF factors (C-D), color-coded by treatment days and cell states.

### SUPPLEMENTARY TABLE LEGEND

**Supplementary Table 1.** Excel sheet containing the top 100 genes in each factor for the scHPF model of the *in vitro* time course.
